# *Nanos2+* cells give rise to germline and somatic lineages in the sea anemone *Nematostella vectensis*

**DOI:** 10.1101/2023.12.07.570436

**Authors:** Andreas Denner, Julia Steger, Alexander Ries, Elizaveta Morozova-Link, Josefine Ritter, Franziska Haas, Alison G. Cole, Ulrich Technau

**Affiliations:** Department of Neurosciences and Developmental Biology, Faculty of Life Sciences, University of Vienna, Djerassiplatz 1, 1030 Vienna, Austria; Research platform SINCEREST, University of Vienna, Djerassiplatz 1, 1030 Vienna, Austria; Max Perutz labs, University of Vienna, Dr. Bohrgasse 7, 1030 Vienna, Austria

## Abstract

In all animals, stem cell populations of varying potency facilitate regeneration and tissue homeostasis. Notably, germline stem cells in both vertebrates and invertebrates express highly conserved RNA-binding proteins, such as *nanos, vasa* and *piwi*. Interestingly, in animals, which are capable of whole-body regeneration, such as poriferans, hydrozoans and planarians, these genes are also expressed in somatic multi- and pluripotent stem cells, which led to the proposal that they had an ancestral role in all stem cells. While multi- and pluripotent interstitial stem cells have been identified in hydrozoans, they have not unambiguously been demonstrated in other cnidarian classes. Therefore, it is currently unclear if these stem cell systems share a common evolutionary origin or have been adapted individually in different lineages as homoplasy. We therefore aimed to characterize stem cells expressing conserved stem cell marker genes in the sea anemone *Nematostella vectensis*, to gain insight of shared traits governing the regulation of this enigmatic cell type. Through single cell transcriptomics, we identify cell populations expressing the germline associated markers *piwi1* and *nanos2* in the soma and germline. Transgenic reporter genes reveal a lineage giving rise to somatic cells, consistent with a role as a multipotent stem cell population. Cell proliferation studies show that a fraction of *nanos2+* reporter cells are cycling and CRISPR/Cas9 mediated gene knockout show that *nanos2+* progenitor cells are indispensable for male and female germline maintenance in *Nematostella*. This suggests *nanos* and *piwi* genes have a conserved role in somatic and germline stem cells in cnidarians.

## Introduction

Cnidarians are well known for their high regenerative potential, sexual as well as asexual reproduction and the potential for extreme longevity with some species being even considered immortal (1–6). Other metazoans, which share these features, facilitate them through multi- or pluripotent stem cell populations like the archaeocytes of sponges or the neoblasts of planarians (7,8). However, it is currently unclear if the diverse stem cell systems observed in the animal kingdom are homologous. The comparison of stem cells in bilaterians with those of basally branching animals, such as cnidarians, may shed light into their potential common ancestry. The freshwater polyp *Hydra* possesses four independent stem cell populations, the ectodermal and endodermal epithelial stem cells, as well as two sub- populations of interstitial or i-cells, one that gives rise to gametes, and one that gives rise to gland, mucous, nematocytes and neuronal cells (9–13). Yet, i-cells have so far only been observed in hydrozoans, where they can be easily identified morphologically and by a high nucleus to cytoplasm ratio. For the rest of the cnidarian phylum, the source of new cells (i.e., their stem cells) for any tissue is unknown. While in several non-hydrozoans, “amoebocytes” have been described, their differentiation status and relationship to interstitial cells is unclear (10). The lack of clear evidence of i-cells outside of hydrozoans suggests that this cell type may be an innovation of this lineage rather than a synapomorphy of the Cnidaria as a whole (10). This would be somewhat surprising given that most non-hydrozoan cnidarians have similar growth and regeneration capacities as hydrozoans. Hence, if there are no i-cells outside of hydrozoans, the question is, which cells can fulfill their function. Alternatively, they might have been simply overlooked by the so far classic histological methods used to address this question. Therefore, we aimed to elucidate the characteristics of the ancestral state for application to evolutionary questions and deduce common principles of cnidarian stem cells, by utilizing the anthozoan *Nematostella vectensis* as a model system. Since in *Nematostella* no cell population similar to the hydrozoan i-cells has been described, we performed a molecular characterization of potential stem cells through the expression of stemness-associated marker genes (14). As an alternative way to the identification of stem cells by morphological features, we reasoned that conserved molecular markers of stem cells might point us to cell populations with stem cell properties. Indeed, *piwi, vasa* and *nanos* were found to be expressed in multipotent and pluripotent stem cells of various metazoans (10,15,16). As members of the so-called germline multipotency program (GMP), these genes are often expressed in, or even restricted to the germline of ecdysozoan and vertebrate animals, although they can also be expressed in multipotent stem- and progenitor cells in echinoderms, lophotrochozoans, cnidarians and poriferans (17–19). *Piwi* genes interact with piRNAs, which guide them to repress transposons, and protect the genome from damage (20). The *nanos* gene family codes for zinc-finger containing RNA-binding proteins, with apoptosis and differentiation promoting mRNA targets, thereby inhibiting those processes (21,22). The activity of both gene families is especially conserved in the germline of vertebrates and insects, where they ensure genomic integrity and keep gametes in a stem cell-like state (23–26). In *Nematostella*, we found three piwi genes (one of which appears to be a pseudogene) (27) and two *nanos* paralogs, *nanos1* and *nanos2*. While *nanos1* marks the neuronoal progenitor cells (14), *nanos2* expression studies implied a role in germline formation, similar to that of *vasa1*, *vasa2* and *PL10* (28). Further, *nanos* gene expression can be detected in the interstitial stem cells of *Hydractinia* and *Hydra* (29,30). Together, these expression studies of *nanos* suggested a role in germline, multipotent stem cells and neuronal progenitor cells.

In this study, we used single cell transcriptome data to identify a population of *piwi1* and *nanos2* positive cells. We generated transgenic reporter lines using *piwi1* and *nanos2* promoters and showed that both cell populations give rise to somatic cell types of the neuroglandular lineages. A *nanos2* knockout mutant reveals a conserved crucial role for *nanos* in the formation of the germline.

## Results

### *Piwi1* and *Nanos2* are expressed in a broad range of somatic cell types and gametes

*Piwi1* has been previously shown to be expressed broadly during early development (27), however, expression at later stages has not yet been well described and the fate of the *Piwi1* expressing cells is still unclear. To visualize *piwi1* expression *in vivo*, we generated a transgenic reporter line expressing mOrange2 (mOr) under the control of the *piwi1* promoter. The fluorescent reporter recapitulates the mRNA expression observed by *in situ* hybridization (18; S1 Fig) shifting from a ubiquitous expression in the gastrula stage to mainly being restricted to endomesodermal tissues during polyp metamorphosis. A low level of reporter expression in ectodermal body wall and tentacle epithelia is, however, retained from the polyp stage onwards (Fig 1B,F). In the adult animal, the endodermal tissue surrounding the pharynx and the gonadal region of the mesentery and ciliated tract are also mOrange positive (Fig. 1C). In juvenile animals, reporter expression is more restricted to few cells residing between the distal tip of the retractor muscle and the ciliated tract, the region where the gonad will form during sexual maturation (Fig. 1D, arrowheads). In the adult mesenteries the endomesodermal expression extends to all epithelia surrounding the gonads and the reticulate tract (Fig. 1E). Notably, large numbers of small single cells displaying mOrange reporter expression are found all along the mesenteries, but especially concentrated around the parietal muscle (Fig. 1F,G).

**Fig. 1:**
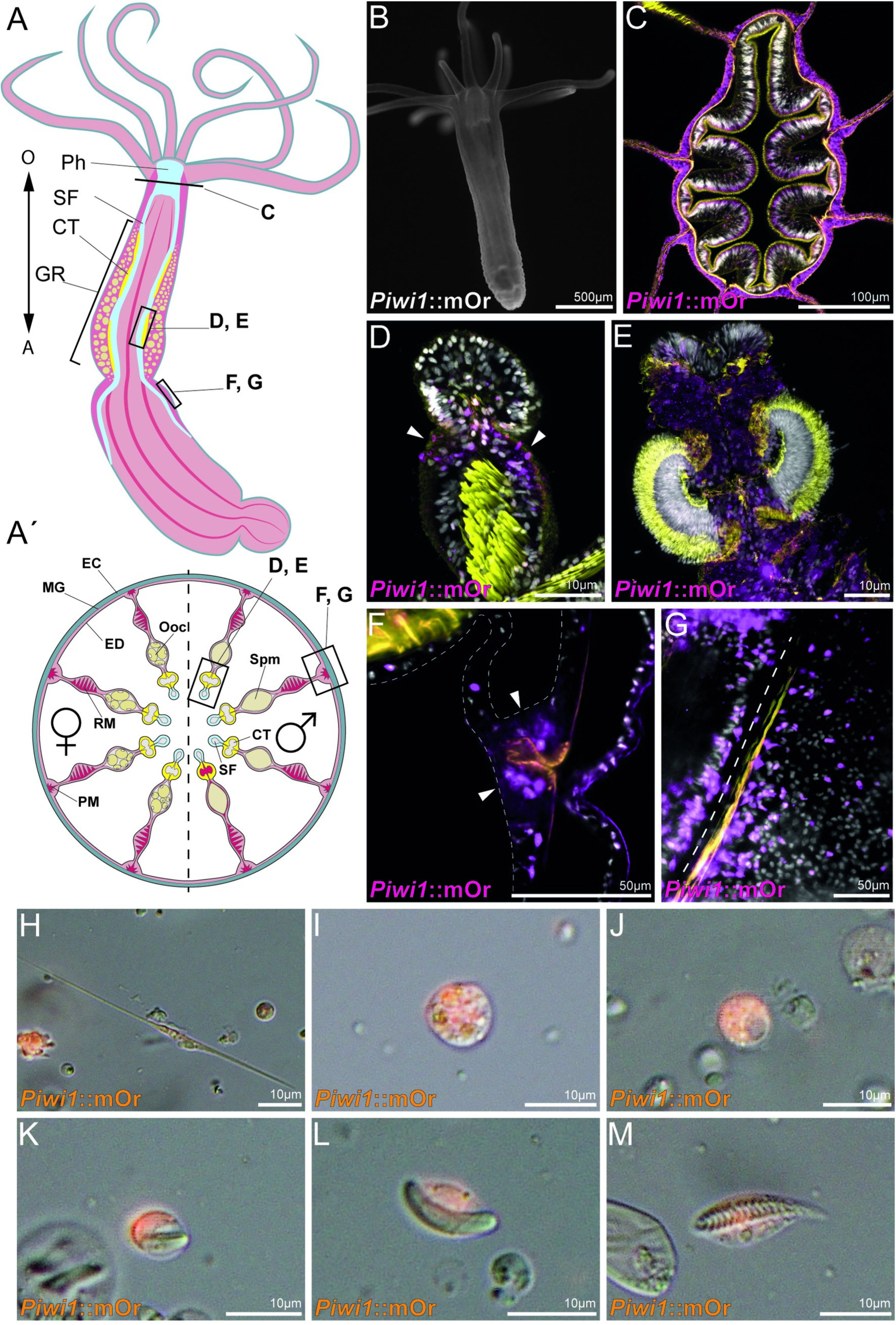
Piwi1 reporter expression in somatic cell types. **A, Á**: Whole-body and transversal section schematic indicating tissues and location of piwi expressing cells. **A**: O: Oral pole; A: Aboral pole; Ph: Pharynx; SF: Septal filament; CT: Ciliated tract; GR: Gonad region; **Á**: EC: Ectoderm; MG: Mesoglea; ED: Endomesoderm; PM: Parietal muscle; RM: Retractor muscle; SF: Septal filament; CT: Ciliated tract; Ooc: Oocytes; Spm: Spermaries. **B**: Juvenile Piwi1::mOr polyp. mOrange is displayed as white. **C**: Section through a pharynx. **D**: Juvenile mesentery; Arrows mark piwi1+ cells accumulating at the gonad primordium. **E**: Ciliated tract of an adult mesentery. **F**: Piwi1+ cells concentrated at the mesentery base, surrounding the parietal muscle (orange). **G**: Optical section along the mesentery base showing piwi1+ cells along the parietal muscle; the dotted line indicates the mesentery base. **H**: Piwi1+ neuron. **I**: Piwi1+ gland cell. **J**: Piwi1+ oocyte. **K**: Piwi1+ immature nematocyte. **L**: Piwi1+ mature nematocyte. **M**: Piwi1+ spirocyte. For all fluorescent pictures DAPI is displayed as white and phalloidin is displayed as yellow.

To identify cell types expressing *piwi1*::mOr, we dissociated adult animals and surveyed single cells. In these cell spreads, we find few *piwi1*::mOrange+ cells in oocytes (Fig. 1J) as well as all descendants of the neuroglandular lineage: neurons (Fig. 1H), gland cells (Fig. 1I) and cnidocytes (Fig. 1K-M). In summary, *piwi1::*mOrange is expressed in oocytes, spermatogonia, putative germline stem cells, cells of the neuroglandular lineage and at low levels in the ectodermal body wall epithelia as well as the endomesodermal epithelia surrounding the gonad region and the pharynx. This situation is similar to hydrozoan cnidarians, where *piwi* expression has been detected in somatic as well as germline cell types (31–33).

We next wished to study the lineage derivatives of *nanos* expressing cells. A phylogenetic analysis shows that the *nanos* gene has duplicated in the common ancestor of all cnidarians and most cnidarians have retained both paralogs (S2 Fig). *Nanos1* has recently been found to mark the neural precursor cells in *Nematostella* (14). To monitor the expression and fate of *nanos2*-expressing cells in *Nematostella*, we generated a transgenic reporter line using the upstream and downstream cis-regulatory elements expressing mOrange. The fluorophore expression during developmental stages recapitulates the pattern retrieved from in situ hybridisation (ISH), changing from an ubiquitous expression in the gastrula and planula stage to being mostly restricted to the primary mesenteries (20, S1 Fig). In juvenile transgenic animals, *nanos2* is expressed around the mouth, similar to hydra (30) in the first forming ciliated tracts of the primary mesenteries, as well as in single cells distributed throughout the whole animal in both germ layers (Fig. 2B). The ectodermal layer displays a dense net of neurons, expressing the mOr fluorophore at low levels (Fig. 2D), whereas in the endomesodermal layer sensory neurons, gland cells and undifferentiated cells are concentrated along the parietal muscle (Fig. 2E). Interestingly, the pharynx shows patchy expression in the ectodermal tissue (which, according to recent profiling has an endodermal identity (34)) with parts of the innermost folds being transgenic as well as the outward folds bordering the inner layer (Fig. 2C). Similar to the *piwi1* transgenic line, *nanos2::*mOrange is expressed in the epithelia surrounding the gonad, however *nanos2*::mOrange additionally marks the side of the ciliated tract bordering the gonadal regions, corresponding to the reticulate tract (Fig. 2F,G). In the single cell spreads we can again detect *nanos2*::mOrange expression in all neuroglandular cell types (Fig. 2H-M). However, in both *nanos2*::mOrange and *piwi1*::mOrange transgenic animals, only a small fraction of neuroglandular cell types show low levels of residual fluorophore, and most are not expressing mOrange. Therefore, we conclude that both genes are most likely expressed in the precursor population, from which the neuroglandular cell types differentiate and carry on residual mOrange reporter protein.

**Fig. 2:**
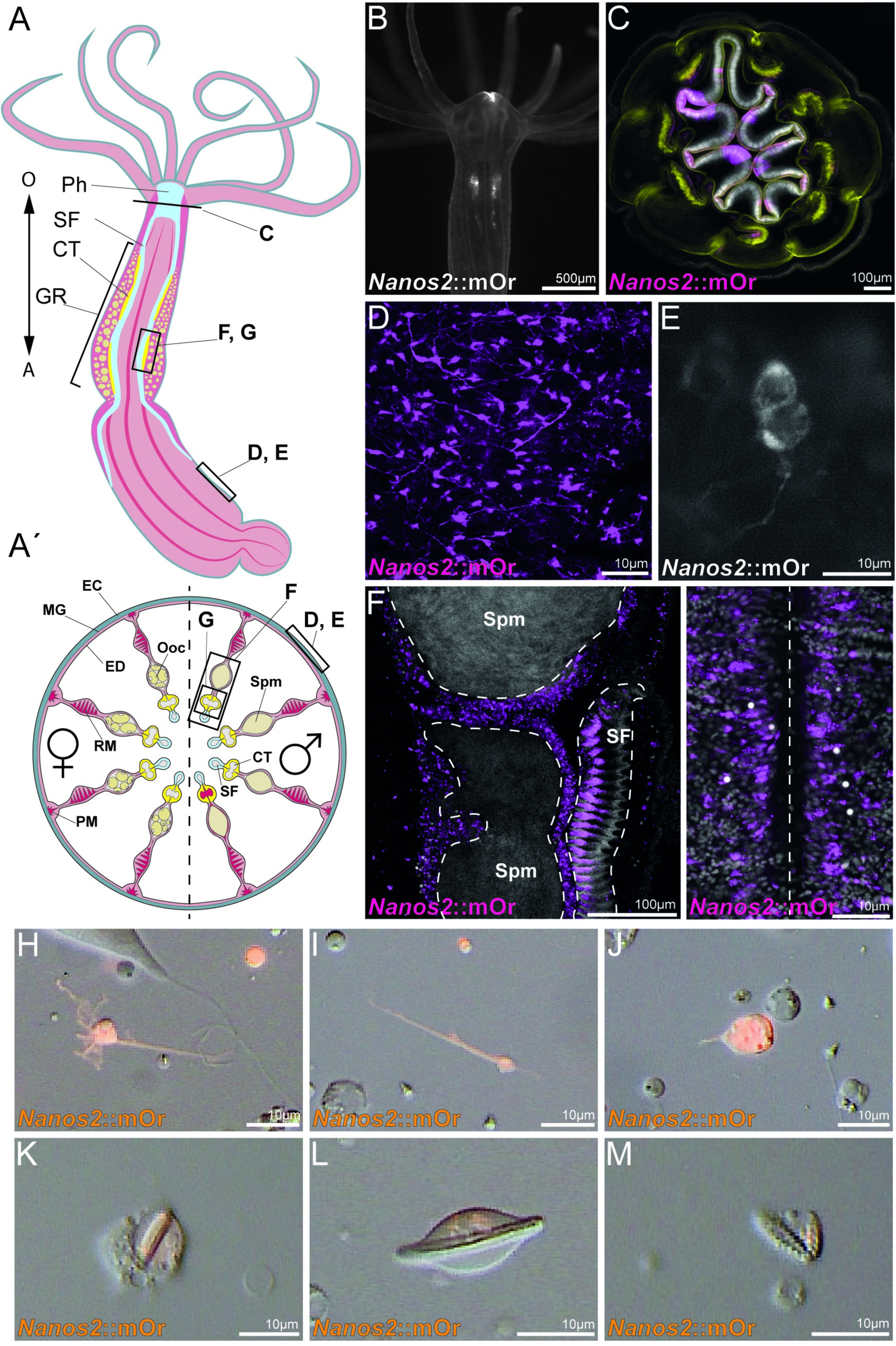
Nanos2 reporter expression in somatic cell types. **A, Á**: Whole-body and transversal section schematic indicating tissues and location of *nanos2::*mOrange expressing cells. **A**: O: Oral pole; A: Aboral pole; Ph: Pharynx; SF: Septal filament; CT: Ciliated tract; GR: Gonad region; **Á**: EC: Ectoderm; MG: Mesoglea; ED: Endomesoderm; PM: Parietal muscle; RM: Retractor muscle; SF: Septal filament; CT: Ciliated tract; Ooc: Oocytes; Spm: Spermaries. **B**: Juvenile *nanos2*::mOr polyp. mOrange is displayed as white. **C**: Section through a pharynx. **D**: neuronal net in the body wall of an adult polyp expressing *nanos2* reporter. **E**: *nanos2* reporter positive cell doublet in the body wall ectoderm. **F**: Vertical section through a male gonad and ciliated tract. The epithelia surrounding the spermaries (Spm) and the bordering septal filament (SF) express *nanos2* mOr. **G**: Optical section along the mesentery base showing *nanos2* mOr positive cells along the parietal muscle; the dotted line indicates the mesentery base. H-M: *nanos2* mOr positive cell types. **H**, **I**: neurons. **J**: gland cell. **K**: immature nematocyte. **L**: mature nematocyte. **M**: spirocyte. For all fluorescent pictures DAPI is displayed as white and phalloidin is displayed as yellow.

Taken together, both *nanos2* and *piwi1* reporter lines show a very similar expression pattern in terms of cell types. The *nanos2* promoter drives fluorophore expression in all descendants of the neuroglandular lineage and the gonad epithelia. However, in contrast to *piwi1*::mOrange, *nanos2::*mOrange is not expressed in oocytes, but marks the epithelia around the mouth, in the pharynx and the putative gametogenic region of the reticulate tract in the septal filament (35).

In contrast to most bilaterian animals, gametes in cnidarians are presumably not formed by a set-aside cell population, but rather constantly produced by somatic tissue (36). This suggests the presence of a population of gametogenic stem cells. As both *nanos* and *piwi* genes are conserved germline factors, we studied their expression in detail in the gonadal region. Both genes show the same expression pattern in males where they are expressed in the outer perimeter of the spermaries (Fig. 3C,E). In females, however, *nanos2* marks the trophocytes (accessory cells), which are situated close to the germinal vesicle of each oocyte (Fig. 3B,F). These cells form the trophonemata, a cnidarian specific ring-shaped structure consisting of multiple trophocytes, which are in close contact with the surface of the oocyte and thought to provide nutrients to the growing egg (Fig. 3G, Arrowhead; (37,38)). Interestingly, a plug of ciliated cells, which express both *nanos2* and *piwi1* is apparent at each sperm cyst. This structure has been described previously, and has been speculated to be homologous to the trophonemata (Fig. 3H,I, arrowheads; (39)).

**Fig. 3:**
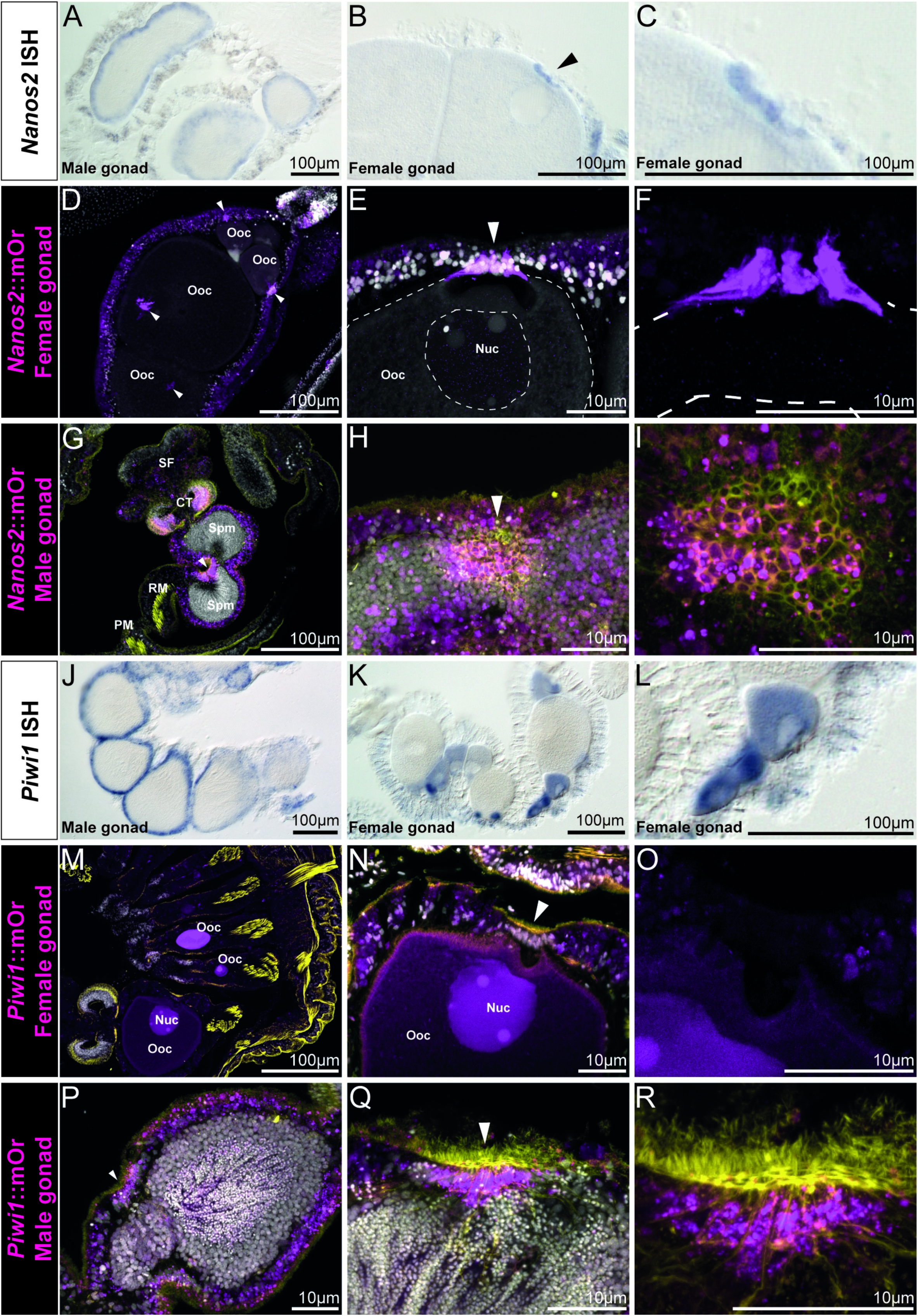
Nanos2 and *piwi1* mRNA and reporter expression in gametes and the gonadal region. **A**: *Nanos2* expression in male mesenteries. **B**: *Nanos2* expression in female mesenteries. *Nanos2+* trophocytes are marked by an arrow. **C**: Detail picture of two *nanos2::*mOr expressing trophocytes in B. **D-I**: *Nanos2*::mOr expression in adult gonads. **D**: Female gonad showing reporter expressing trophonemata located closely to the germinal vesicle of each oocyte are indicated by arrowheads. **E**: *Nanos2*::mOr expressing Trophonema (arrowhead), just above the pronucleus of the oocyte. Nuc: Nucleus of the oocyte or germinal vesicle. **F**: Zoom picture of the trophocyte cells comprising the trophonema in E (DAPI is not displayed in this picture). **G:** Male gonad, with mOr expressing in the gastrodermis around the spermaries and the bordering tissue of the ciliated tract. The ciliated plug is indicated by an arrowhead. **H**: Detail picture of a ciliated plug (arrowhead) in the gastrodermis surrounding the sperm chambers. Cells composing the plug can be identified by increased mOrange expression and phalloidin incorporation (yellow). **I**: Zoom picture of the ciliated plug in H (DAPI is not displayed in this picture). **J**: *Piwi1*::mOr expression in male mesenteries. **K**: *Piwi1*::mOr expression in female mesenteries. **L**: Zoom picture of the mOr+ maturing oocytes in K. **M-R**: *Piwi1*::mOr expression in adult gonads. **M:** Female gonad. The mesentery is folded multiple times and three mOr+ oocytes (Ooc) are visible. **N**: Detail picture of an oocyte. The mOr- trophonema is indicated by an arrowhead. **O**: Zoom picture of the trophonema in N (DAPI is not displayed in this picture). **P**: Male gonad expressing *piwi1*::mOr in the endomesoderm around the sperm chambers and the bordering tissue of the ciliated tract. The ciliated plug is indicated by an arrowhead. **Q**: Detail picture of a ciliated plug (arrowhead) in the gastrodermis surrounding the sperm chambers. Cells composing the plug can be identified by increased mOrange expression and phalloidin incorporation (yellow). **R**: Zoom picture of the ciliated plug in Q (DAPI is not displayed in this picture). For all fluorescent pictures mOr is displayed as magenta, DAPI is displayed as white and phalloidin is displayed as yellow. **Legend**: EC: Ectoderm; MG: Mesoglea; ED: Endomesoderm; PM: Parietal muscle; RM: Retractor muscle; GR: Gonadal region; Ooc: Oocytes; Spm: Spermaries; CT: Ciliated tract; SF: Septal filament.

*Piwi1* is expressed in the oocytes itself, where the reporter concentration is highest in smaller oocytes and fading in larger ones (Fig. 3J). However, *piwi1* expression is lacking from the trophonemata (Fig. 3K) but can also be detected in the ciliated plugs of the spermaries (Fig. 3L, M, arrowheads). To our knowledge, this plug has not been functionally described in *Nematostella* and we can only speculate about its function. It is apparent that the size of the nuclei in the sperm cysts decreases from the outer margins towards the inside, suggesting that sperm differentiation proceeds from the perimeter inwards. These small nuclei, however, seem to flow towards the ciliated plugs, which indicates that this might be the place where sperm is released into the body column during spawning (Fig. 3H, L, M).

Through the expression patterns obtained by ISH and the reporter gene expression in transgenic animals, we conclude that both *piwi1* and *nanos2* are expressed in somatic cell types of the neuroglandular lineage and the germline. The mOr reporter of both genes is detected only in a few cells of each of the somatic cell types. This suggests that the fluorophore in these fully differentiated cells was carried over from a common NPC. However, *piwi1* is expressed much stronger in the germline, whereas *nanos2*-positive cells are much more frequent in the descendants of somatic lineages. Also, both markers are expressed in epithelial cell populations, where *piwi1* is very broadly expressed in the body wall ectoderm at a low level, while *nanos2* is restricted to regions of the pharynx and ciliated tract, with a higher expression level overall. For evaluation of functional studies affecting the expressing cell populations, we therefore focused on surveying the *nanos2* expressing population.

### *Nanos2+* cells are a cycling population

To assess if the *nanos2*::mOrange expressing cells comprise an actively cycling population, we labeled transgenic animals for 24 hours with EdU. After this period, a fraction of the mOrange+ cells in each body region and tissue were marked by EdU (Fig. 4). Since the transgenic line retains mOrange reporter expression in derivatives of *nanos2* expressing cells, we suspect, that the EdU-negative *nanos2*::mOrange+ cells is comprised of terminally differentiated cell types such as neurons and gland cells as well as possibly slow cycling quiescent stem cells, whereas the fast cycling cells may be immediate precursor populations of the neuroglandular or germline cells.

**Fig. 4:**
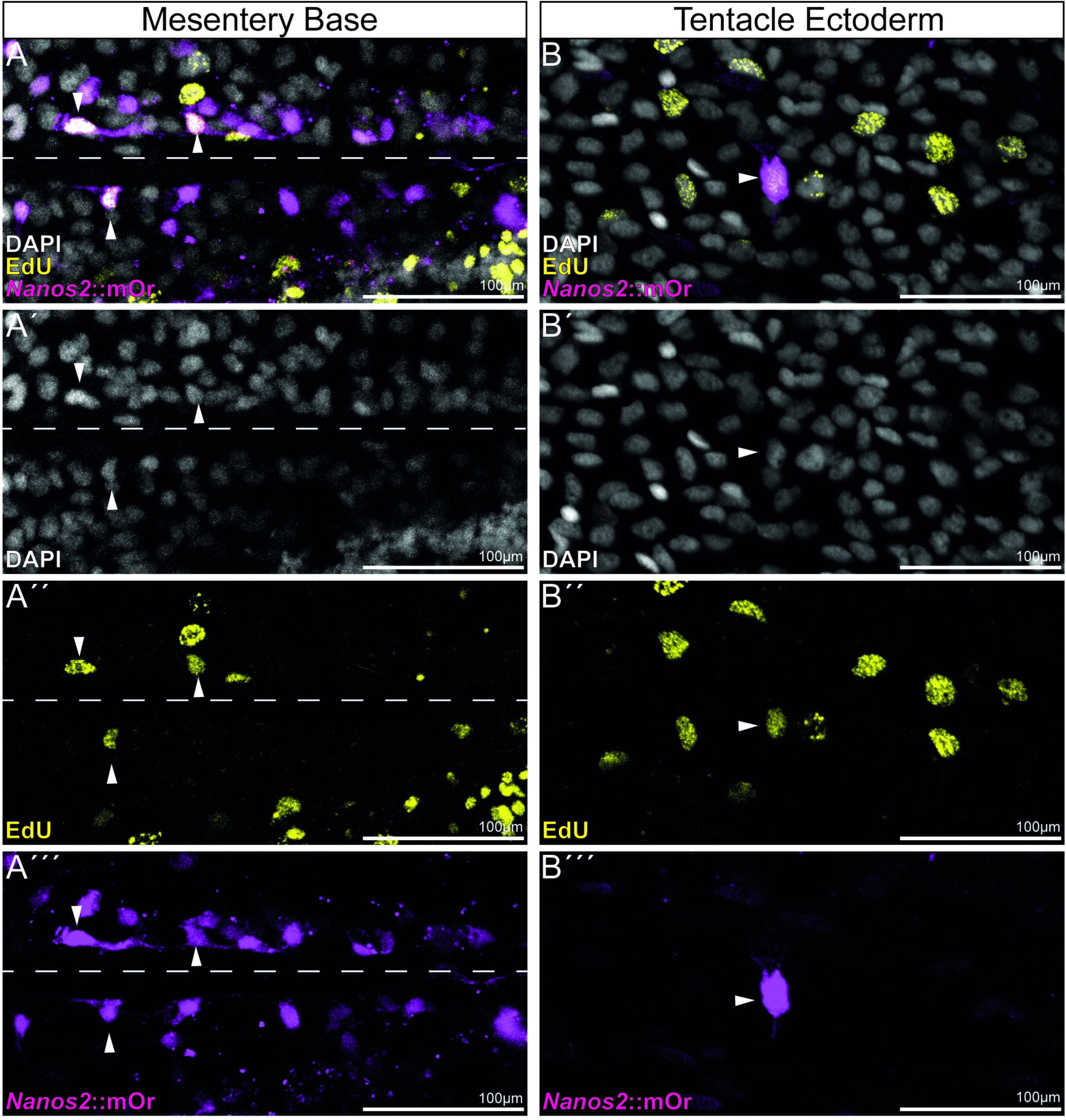
EdU incorporation of *nanos2*::mOrange expressing cells. **A-Á’’**: Fluorescent picture of the endoderm at the base of the mesentery. The attachment site of the mesentery is indicated by a dotted line. mOrange / EdU+ cells are indicated by arrowheads. **B-B’’’**: Fluorescent picture of the tentacle ectoderm. mOrange / EdU+ cell is indicated by an arrowhead.

To determine this fraction of actively cycling *nanos2* expressing cells, we wished to carry out a double staining of *nanos2* mRNA and EdU pulse labeling. However, the cellular resolution of the *in situ* hybridization of *nanos2* in adults is very limited, also because the density of small *nanos2* expressing cells is so high in many regions of the body. Therefore, we subjected the transgenic animals to a EdU pulse of 5 hours, dissociated them afterwards, and counted the numbers of EdU positive and negative cells as well as *nanos2*::mOrange positive and negative cells (Supporting Information, Table 1). We found that *nanos2*::mOrange cells comprise a significant fraction of all cells (23%), of which 11.8% were EdU+ under these conditions. This shows that at least a subpopulation of *nanos2* positive cells are proliferative. Together with our finding that the mOrange reporter protein is expressed in somatic lineages this is consistent with the possibility that *nanos2*-expressing cells have properties of multipotent stem cells or progenitors. However, we also note that among the *nanos2*::mOrange negative cells, a similar fraction (13,3%) is also labeled by EdU. The nuclei of these *nanos2*-negative/EdU+ cells are relatively large and round, raising the possibility that these might be epithelial cells. While the nature of these cycling cells remains elusive, this shows that *nanos2*-expressing cells are not the only cycling cells in an adult polyp.

Some adult stem cells are mitotically quiescent for long times and rarely divide. These can only be detected by a label retention assay, where an EdU pulse is followed by long periods of no label. Under such conditions, rapidly cycling progenitors would rapidly dilute out the incorporated EdU label to the progeny cells, while quiescent or rarely dividing stem cells would retain the label even after an extended time. To test whether *Nematostella* might have slowly or quiescent stem cells, we incubated the *nanos2*::mOrange (Fig. 5) and *piwi1*::mOrange transgenic polyps (S3 Fig) for 1 week in EdU, followed by extensive chase with *Nematostella* medium and fixed the animals after several time points up to 3 months after the labeling. Figure 5 shows that in cryosections of the polyps we found indeed several EdU^+^/*nanos2*:mOrange^+^ cells 1-3 months after the pulse, demonstrating the existence of quiescent or rarely dividing stem cells. Since the label could be also retained by differentiated, long-lived cells, e.g. neurons, it is important to note that the morphology of these cells were small round cells, intercalated between the epithelial cells throughout the animal, but often in the vicinity of the mesentery base, suggesting that these are non-differentiated cells, similar to interstitial cells in hydrozoans. Thus, the detection of rarely dividing *nanos2*+ cells explains at least in part the relatively low fraction of pulse-labeled cells.

**Fig 5:**
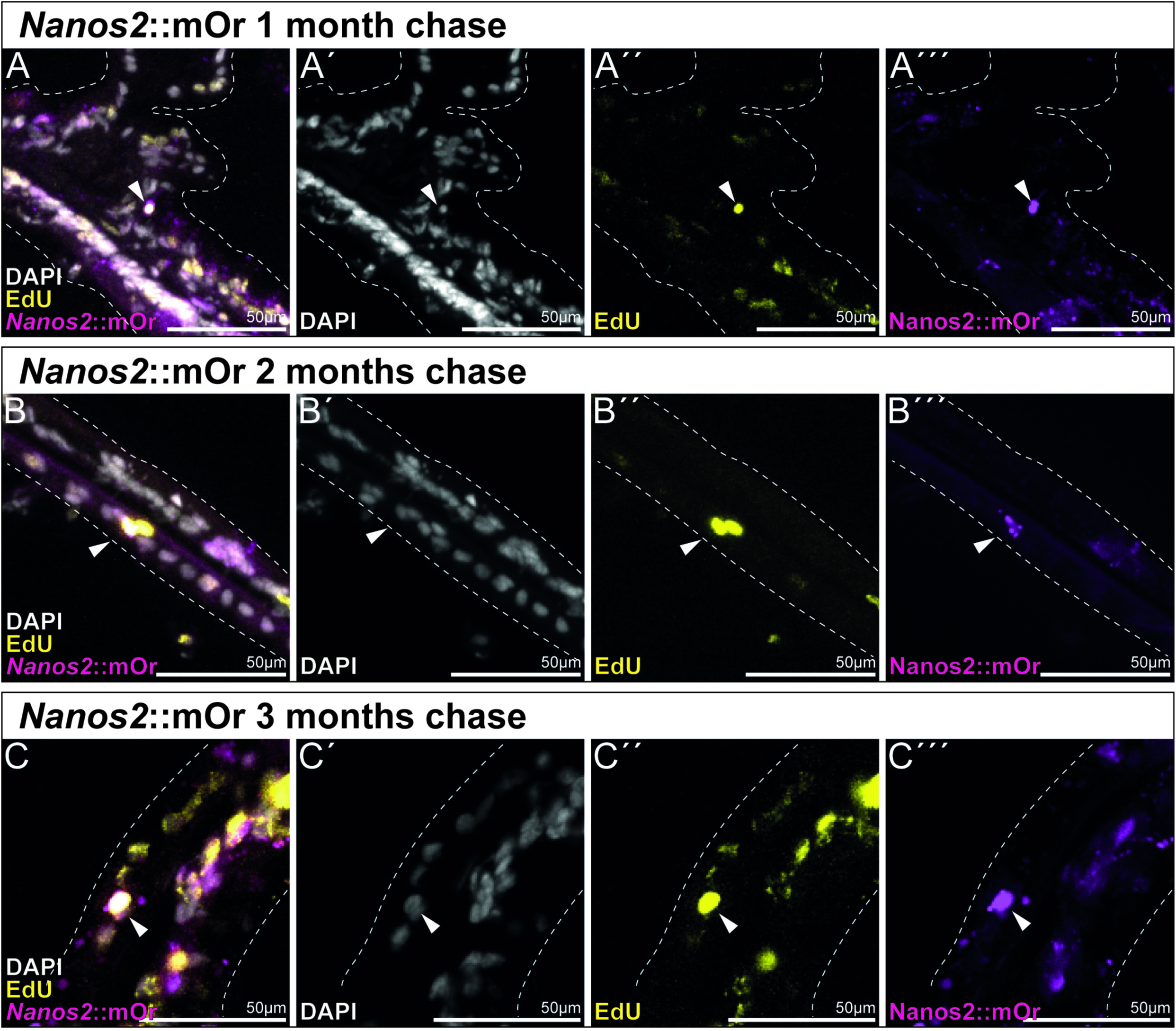
Long term EdU label-retention assay shows that *nanos2*::mOr marks slow cycling, quiescent cells over 3 months. **A-C**: EdU signal is retained in few mOr+ cells in the body wall and mesentery epithelia over 3 months, marking a subpopulation of slowly cycling cells that still proliferate, indicated by EdU+ cell doublets (**B**). **A**: 1 month label retention. **A**’: DAPI only. **A**’’: EdU only. **A**’’’: mOrange only. **B**: 2 month label retention. **B**’: DAPI only. **B**’’: EdU only. **B**’’’: mOrange only. **C**: 3 month label retention. **C**’: DAPI only. **C**’’: EdU only. **C**’’’: mOrange only.

### *Nanos2* mutants fail to form primordial germ cells and gametes

To test the function of *nanos2* in cellular differentiation, we generated a KO mutant using CRISPR/Cas9 (40), introducing a 1bp insertion at the 5’end of the predicted Nanos specific zinc finger sequence (S4 Fig). This results in an early stop codon, which renders the protein unable to form a ternary complex with binding partners such as Pumilio or bind to mRNAs on its own (41,42). After crossing the heterozygous mutant F1 parent generation, we observed no increased mortality or developmental defects in the offspring, and homozygous mutant animals were only identifiable through sequencing of the *nanos2* locus. We raised the homozygous mutant polyps to adulthood and tried to spawn them. Yet, while wildtype polyps start to spawn after about 4-5 months postfertilization, even after 1 year none of the mutants spawned any gametes, neither oocytes nor sperm, suggesting that the *nanos2* mutants are sterile.

To further validate this, we carried out a staining for Vasa2 protein. Vasa has been established as a marker for the germline in *Nematostella*, *Hydra* and several other animals (17,27). In *Nematostella*, Vasa2 protein has been shown to mark putative germline stem cells during development, spermatogonia, oocytes and single cells in the epithelium surrounding the gonads (27). In line with that, in wild type animals, Vasa2+ PGCs are visible between the septal filament and the gonad, closely associated with the reticulate tract, a region of the ciliated septal filament most closely bordering the gonad (Fig. 6C-F,K). *nanos2*-/- animals on the other hand, display somatic gonads lacking oocytes or sperm and devoid of Vasa2+ cells (Fig. 6G,H). This result underlines the observed sterility of adult mutants.

**Fig. 6:**
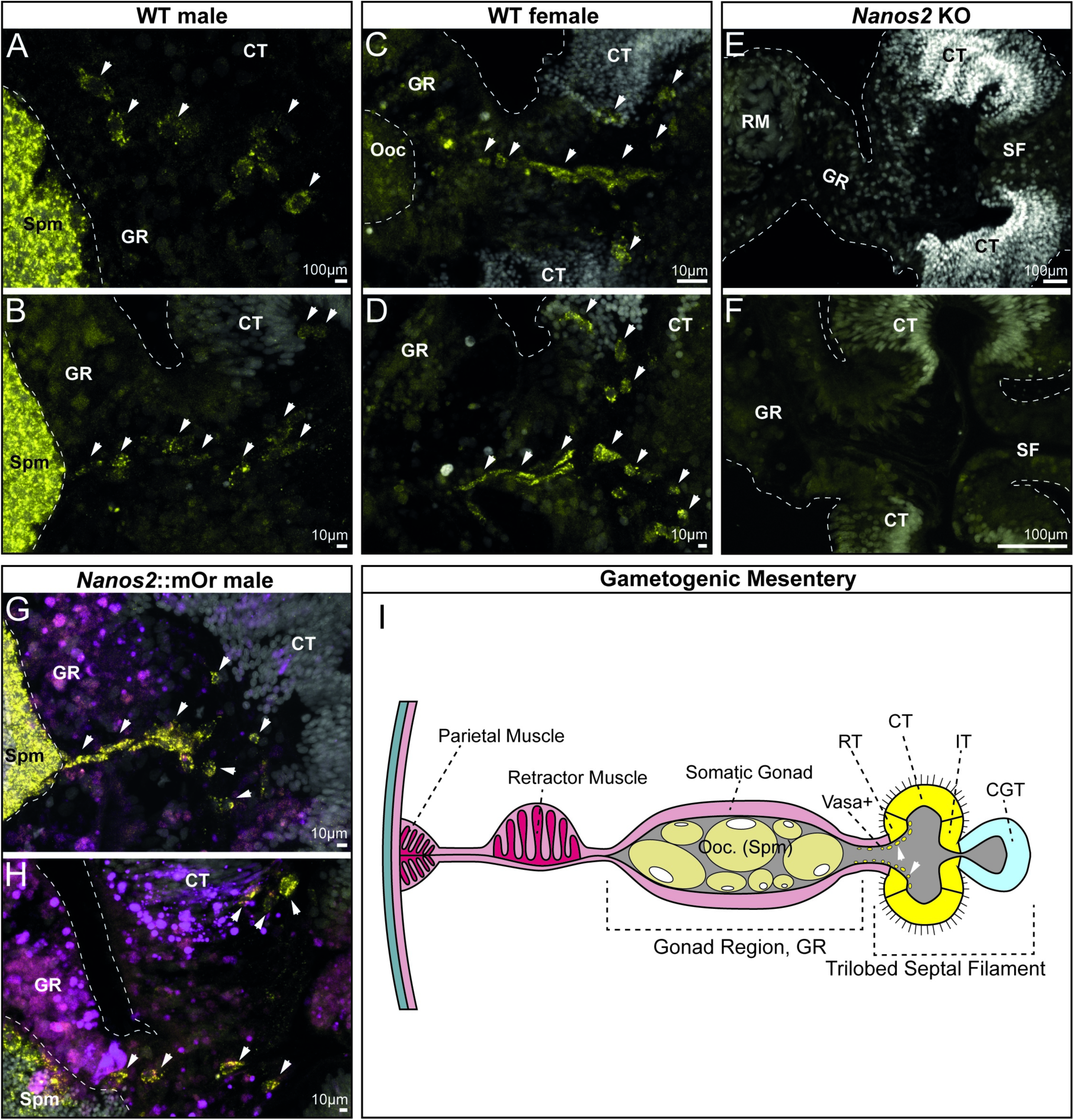
Vasa2 immunostaining of PGCs confirms the absence of germline cells in *nanos2*-/- mutants. **A-H:** Transversal sections of the gonad region and trilobed septal filament of WT male (**A, B**), WT female (**C, D**), *Nanos2* KO (**E, F**) and *Nanos2*::mOr (**G, H**; mOr fluorophore is depicted in magenta) animals, stained with DAPI (white) and Vasa2 antibody (yellow). Vasa2+ cells have been indicated by white arrows. Note, that no Vasa2+ positive cells nor mature oocytes or spermatocytes could be detected in *Nanos2* mutants, explaining the sterility of the animals after 12 months. If not otherwise indicated, scale bars correspond to 10µm. **I:** Schematic of a gametogenic female mesentery with a trilobed septal filament. vas+: Vasa2 positive putative primordial germ cells; ret. t.: Reticulate tract; cil. t.: Ciliated tract; int. t.: Intermediate tract; cni.-gla. t.: Cnido-glandular tract.

### Nanos2 is required for germline formation but dispensable in somatic cell type lineages

The characterization of our *nanos2::mOrange* and *piwi1*::mOrange transgenic reporter lines revealed fluorophore expression in a variety of somatic cell types in both cell layers (see Figs. 1-3). To assess the consequences of the *nanos2*-/- mutation on the cell type composition of the animal, we carried out single-cell RNA sequencing of an individual *nanos2*-/- mutant 2-month-old juvenile polyp. We integrated the *nanos2*-/- mutant cell transcriptomes with the available wildtype adult tissue catalog (14,43), newly mapped to the updated genome and transcriptome models (44). We find that the *nanos2*-/- mutant exhibits no differences in the overall cell type diversity (Fig. 7A-C), further supporting the observation that the mutant is vital and shows no apparent deficiencies of somatic tissues.

**Fig. 7:**
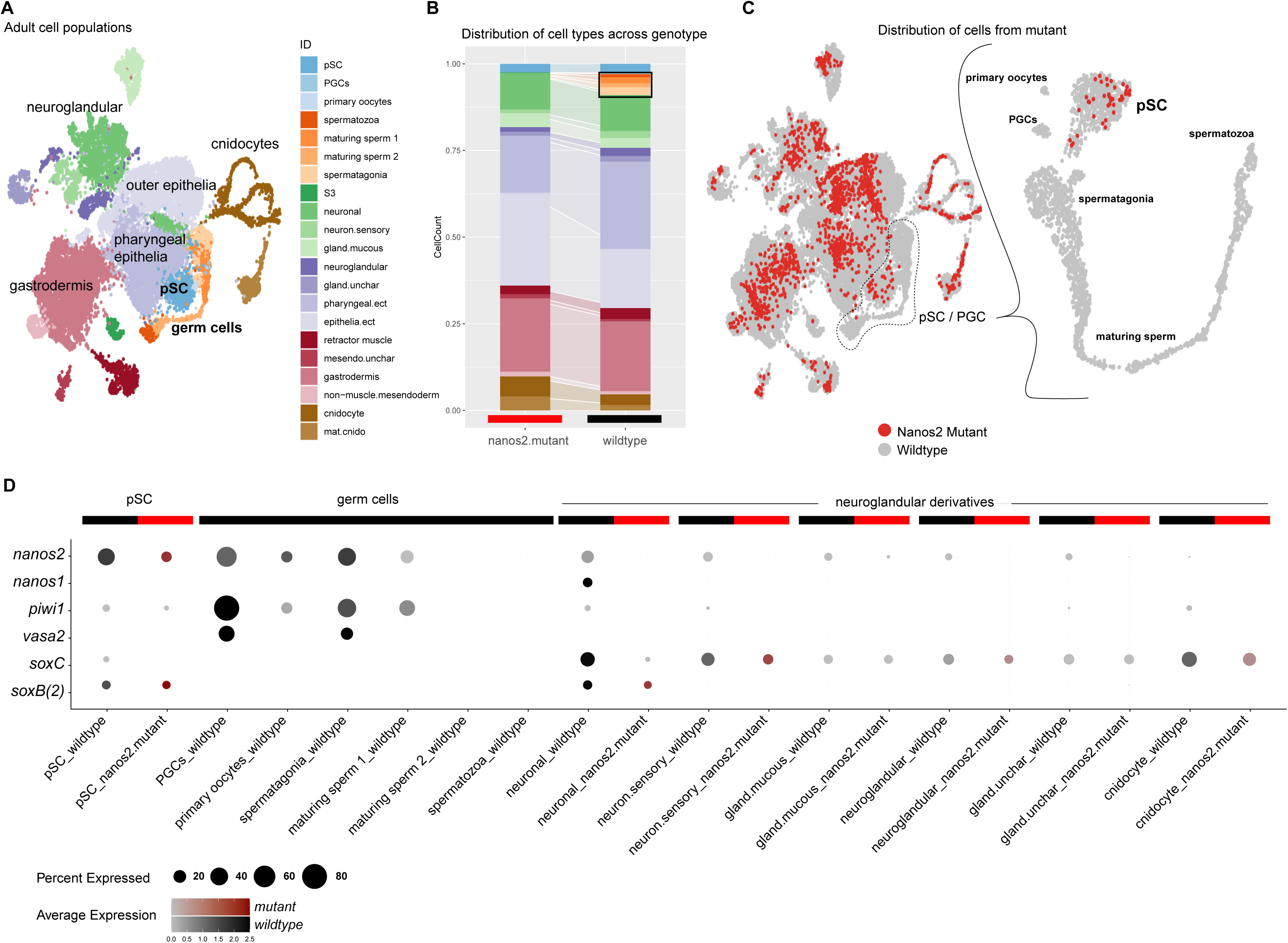
Single cell RNAseq analysis of the *nanos2*-/- mutant reveals absence of primordial germ cells and gametes. **A)** Single-cell transcriptomic map of an individual *nanos2*-/- mutant juvenile polyp merged with wild-type adult tissue libraries. **B)** Library contribution to the map in (A) illustrates a similar contribution of the mutant to all principal cell populations with the exception of germ cells and uncharacterized cell type S3 (black outline). **C)** Distribution of mutant cells (red) across the entire dataset (grey), and the cluster of putative stem cells and germ cells identifies the absence of germ cells and differentiated gametes in *nanos2*-/- mutants. **D)** Expression dotplot of putative stem cells (pSC), germ cells, and neuroglandular derivative cell populations. Detection of gene expression in at least 5% of the cell population is illustrated. Germ cell marker expression (*nanos2, piwi1,* and *vasa2*) shows all three genes are co-expressed and elevated within the wildtype germ cell populations, as well as detected in both mutant and wild type putative stem cells. *nanos2* is also broadly detected across all populations in the wild type samples and is partially co- expressed with markers of early neuroglandular progenitors *nanos1, soxC and SoxB*(*2*), whose expression is unaffected in the mutant.

In the wildtype, we detect elevated levels of *nanos2, piwi1, and vasa2* expression in the putative stem cell and primary germ cell (pSC/PGC) cluster (Fig. 7D). Of note, these genes show a partial overlap with the early regulators of the neuroglandular lineages *nanos1, soxC, soxB2a* (also known as *soxB*(*2*)). To investigate the distribution of the mutant-derived cells within this population, we re-analyzed this cluster separately. Primordial germ cells first originate at the boundary of the ectodermal and mesodermal cell layer in the pharynx of primary and juvenile polyps, from where they migrate into the gonad primordia (45). Within this subset of the single cell data of the wildtype polyp we can clearly identify the maturing spermatozoa, primary germ cells, and a cluster of primary oocytes in addition to the putative stem cell population (Fig. 7C). Importantly, we find no *nanos2*-/- mutant cells contributing to the primordial germ cells, although this sample does contribute to the putative stem cell precursor population (Fig. 7C). Neither male nor female gametes could be detected in *nanos2*-/- mutants. This suggests that *nanos2* is required for the formation of the PGCs in both sexes and subsequent differentiation of gametes at an early stage of development. Together with the morphological observations described above, we conclude that *nanos2* is required for germline formation and gamete differentiation in *Nematostella*.

We note that in the single cell transcriptomes of the *nanos2-/-* mutants, the derivatives of the neuroglandular lineages are still present (Fig. 7A-C), although neuroglandular derivatives are found in the *nanos2* transgenic line, suggesting a lineage relationship (Fig. 2). We previously described *nanos1* marking the neuronal progenitor population, overlapping with *soxC*, which acts as the most upstream determinant of several somatic lineages, such as cnidocytes, neurons, and gland cells (14). This is in line with the observation that the mutants are phenotypically normal, can feed and move as wildtype animals. Long-term observations of our animals did not indicate defects in the homeostatic renewal of neuroglandular cell types or a negative impact on the survivability of *nanos2*-/- mutants. Therefore, while *nanos2* is expressed in NPCs, it is unlikely that *nanos2* is essential for the differentiation of the neuroglandular lineages into terminal cell types. However, we cannot rule out that the lack of a phenotype in the neuroglandular lineages in the *nanos2-/-* mutant might be due to genetic compensation, for instance by *nanos1* (46). In line with this, *nanos2* and *nanos1* also show a partially overlapping expression (Fig. 7C).

Co-expression of *nanos2* and *piwi1* in the neural progenitor population and multiple derivatives, together with expression patterns of the fluorophore detected in our transgenic lines, indicate a role of *nanos2* and *piwi1* beyond the specification of primordial germ cells. However, slight differences in gene function might be inferred by the single cell expression data. *Nanos2* appears to feature a much broader expression profile than *piwi1* and is particularly upregulated in the pSC population of both wild type and mutant. On the other hand*, piwi1* and *vasa2* are highly expressed in the gametes rather than the putative stem cells. The lack of a germline in the *nanos2* mutant animals suggests an epistatic relationship, in which *nanos2* might facilitate the differentiation of pSCs towards *piwi1/vasa2+* gametes, although further studies are needed to confirm this hypothesis. Taken together, our data suggests that *nanos2* and *piwi1* function at the top of the differentiation cascade of most cell types, with the possible exception of epithelial cells, and therefore we consider *nanos2* and *piwi1* positive cells as a population of putative stem cells that give rise to both primordial germ cells and neural progenitor cells, reminiscent of the hydrozoan i-cell populations (47).

## Discussion

Cnidarians have a remarkable tissue plasticity and regenerative capacity. They can grow and shrink repeatedly yet maintain a homeostatic ratio of all cell types at all times. However, many cell types, such as neurons, cnidocytes, and gland cells are terminally differentiated. Importantly, gametes can also be considered a differentiated cell type. Therefore, they must be replenished from a pluripotent or multipotent stem cell pool. In *Hydra*, interstitial stem cells have been shown to act as multipotent stem cells for all cells except epithelial cells (49). In the marine hydrozoan *Hydractinia*, interstitial cells have been shown to also give rise to epithelial cells, thus, they are considered as pluripotent cells (50,51), similar to neoblasts in planarians. Yet, interstitial stem cells have so far only been identified in hydrozoans, and not in any of the other cnidarian classes, despite the fact that they have very similar properties. Therefore, we used conserved molecular stem cell marker genes to molecularly identify putative stem cells in the sea anemone *Nematostella*. This includes the *piwi* and *nanos* gene families, which have been implicated in the germline of many bilaterians (41,52–61). Piwi binds to piRNAs, which are silencing transposons and hence is thought to contribute to the genome integrity of the germ cells (26). However, an involvement in somatic cells has also been demonstrated. For instance, in planarians *piwi* was shown to be essential for the pluripotent stem cells, the neoblasts, which give rise to all other cell types and hence are crucial for maintenance of the whole body (62–64). Additionally, the *piwi1*::mOr transgenic line displays low level expression in the ectoderm and gastrodermis. We could also detect mRNA transcripts of *piwi1* and *nanos2* in cells of these epithelia in scRNAseq, while we were unable to detect them by ISH. The low-level expression of these genes might hint towards the existence of epithelial cells as separate stem cell lineages which replenishes the epithelia of the polyp, similar to those of *Hydra*. This would be in line with the existence of a cycling cell population, which is negative for *nanos2*::mOrange and *piwi1*::mOrange.

Nanos, an RNA-binding zinc finger protein implicated in translational repression (65), has also been implicated in the germline of many bilaterians, but also in somatic stem cells of some highly regenerative animals, e.g. in neoblasts in planarians (66), archeocytes in sponges (67) and interstitial stem cells in *Hydra* (30). In *Hydra*, a transgenic reporter line under control of the *nanos1* promoter showed that *nanos1* positive stem cells give rise to cnidocytes and neurons (48). However, while the function of *piwi* and *nanos* has been shown in planarians, in *Hydra* there are only expression studies available. Moreover, interstitial stem cells have only been demonstrated in hydrozoans, not in other cnidarians. Thus, the nature of possible multipotent stem cells remains enigmatic at this point. The expression of two *nanos* paralogs, *nanos1* and *nanos2* (common to all cnidarians) and two *piwi* genes, *piwi1* and *piwi2*, have been reported before (27,28). However, the cellular resolution of *in situ* hybridization did not permit visualization of individual cells in the adult. Moreover, neither their lineage relationship with other cell types nor their function is known. In the sea anemone *Nematostella vectensis*, single cell RNAseq and transgenic lines showed that *nanos1* is expressed in neuronal precursor cells and gives rise to neuronal cells (14). By contrast, single cell RNAseq suggested that *piwi1* and *nanos2* are expressed in a broad spectrum of undifferentiated and differentiating cell types. In this study, we have generated transgenic lines of the *piwi1* and *nanos2* genes in the sea anemone *Nematostella vectensis.* Both reporter lines mimic the mRNA expression and show similar, but not identical patterns: *piwi1* is expressed broadly at a low level in the ectodermal layer but is strongly upregulated in many small round cells in the inner (endomesodermal) cell layer as well as in the early stages of germ cells (oogonia and spermatogonia). *Nanos2* and *piwi1* transgene expression is found in numerous very small cells located between epithelial cells of both layers, spread all over the whole body. Since the fluorophore also transmits into somatic cell types, like neurons and gland cells, we assume that the *nanos2*+ cells give rise to differentiated cells of the neuroglandular lineage. In addition, at least a fraction of the *nanos2*+ cells are proliferating and the label-retention assay over three months demonstrates that some are very rarely dividing cells. Similarly, the *piwi1* transgenic line also harbors slow cycling label retention cells. These features are essential criteria for a multipotent stem cell population similar to the interstitial stem cells in *Hydra*. Interestingly, a transgenic *vasa2* line also showed somatic cell type derivatives such as neurons (68). Co-staining of the *nanos2*::mOrange line with Vasa2 antibodies showed partial overlap between Vasa2 positive cells (presumably primordial germ cells) and *nanos2*+ cells. These data strongly suggest a role of *piwi1*, *nanos2* and *vasa2* in a population of multipotent stem cells.

However, our *nanos2-/-* mutant did not show any detectable effect on the neuroglandular lineage, suggesting that *nanos2* is not essential for this somatic lineage. Yet, we cannot rule out that *nanos1*, which is expressed in the proneuronal progenitor cells (14), compensates for the loss of *nanos2* in the neuroglandular lineages. Future work will have to address this possibility. As no functional data are available for *piwi1* and *vasa2*, we don’t know whether these genes are essential for neuroglandular lineages and whether the identified single cells are truly somatic stem cells.

Regardless of the role of *nanos2* in the somatic cell lineages, our *nanos2* knockout mutant clearly shows that it is indispensable for the generation of primordial germ cells and hence for the formation of male and female gametes. As a result, *nanos2* mutants are sterile. A sterility phenotype has been also observed for *nanos* knockouts in the mouse (57) and *Drosophila* (69). In *Drosophila, C. elegans* and mouse, *nanos* is necessary for germ stem cell self-renewal and it prevents precocious differentiation (70–72). Since we cannot detect primordial germ cells in the *nanos2* mutant in *Nematostella*, we assume that this function is conserved between bilaterians and cnidarians and dates back at least to the common ancestor about 600 million years ago. While this study could not confirm the presence of a pluripotent stem cell population, it has recently been proposed that the *piwi1*+ and Vasa2+ cell population is the source of gametes as well as SoxB(2)+ neurons based on lineage analyses of transgenic lines, similar to our approach (68). Our data support this claim and confirm through the characterization of *nanos2* KO mutants, that the Vasa2+ cell population residing in the septal filament is the source of gametes in *Nematostella*.

In summary, our results highlight the broad use of germline multipotency genes in somatic as well as gametic cell types in evolutionary early branching animals. We confirm the role of *nanos* as an ancestral key determinant of germline establishment in the earliest branching metazoan and likely a crucial regulator of somatic cell types as well.

## Material and Methods

### Nematostella culture

Polyps of *Nematostella vectensis* were cultured in 1,6% artificial sea water (*Nematostella* medium, NM) and fed daily with *Artemia nauplii*. The animals were kept at 18°C in the dark, while spawning was induced by exposure to light and increased temperature (73). After fertilization egg packages were treated with 3% Cysteine in NM to remove the egg jelly. Larvae were kept at 21°C until further usage or until they reached the juvenile stage.

### Generation of single-cell suspensions

Animals were collected in pre-coated Eppendorf tubes and washed multiple times in NM. Cell dissociations for juvenile and adult *Nematostella* polyps were performed using a Papain (3.75mg or 37.5U/ml in PBS, Sigma-P4762) and Collagenase (1.25mg or 1000U/ml in PBS, Sigma-C9407) solution in a 1:1 mix, with DTT added to a final concentration of 3.5mM. Whole juvenile animals and pieces of adult animals were dissociated in 1ml of the Col/Pap dissociation solution, overnight at room temperature. The tissue was then carefully pipetted with a cut pipette tip to dissociate into single cells. The state of the dissociation was regularly checked using samples of the solution under a light microscope. If clumps were no longer visible, the cells were pelleted in a pre-cooled centrifuge for 3min at 0.4 g. A washing step with 1%BSA in PBS was performed to stop dissociation after which the pelleted cells were re-suspended in 1%BSA/PBS on ice. Cell viability and concentration were assayed using a Cellometer X2 (Nexcelom). Only cell suspensions with a minimum viability of 80% were used for single cell sequencing.

### Single-cell imaging

For single cell imaging, animals were dissociated as described above. The cell solution was then applied to superfrost slides (Thermo scientific) on which the desired imaging area was previously marked with a hydrophobic barrier pen (Thermo scientific). The liquid was then taken off using a micropipette and replaced by 4% PFA in PBS for fixation. After 2 min the fixative was taken off the slides and the adhered cells were washed 3 times for 2 min each in PBS. For light microscopy, PBS was then replaced with vectashield (Vector laboratories) and a cover slip was applied before imaging.

### Single-cell RNA sequencing

The single-cell suspension was loaded into the 10X Genomics platform for encapsulation. Sequencing library was prepared according to the standard 10X protocol and sequenced according to manufacturer’s recommendations. Raw sequence data was mapped to the vs.2 chromosome genome assembly and accompanying Nv2 gene models (44) using the 10x genomics cell ranger pipeline, without secondary analysis and force recovery of 10,000 cells per library.

### Single cell transcriptomic analysis

Count matrices for wild type adult libraries and the newly generated mutant sample were imported into R and processed with the R-package Seurat vs4 (74). Data were merged, normalized with Seurats default “LogNormalize” function. The 500 most variable genes from each sample were identified with the “FindVariableGenes” function with default parameters. The combined gene set was scaled with the “ScaleData” function binned by sample origin (‘split.by’ function). Informative principle components were chosen as those with the standard deviation greater than 2, and used as input for the clustering analysis. Dimensional reduction was performed with UMAP using the same PCs as in the clustering. For the sPC/PGC subset, the top 2000 variable genes were calculated, scaled across the entire dataset binned by sample, and then processed further as described above. An R script for generating the analysis and figures is available on our GitHub page (https://github.com/technau/Nanos2).

### Generation of transgenic animals

The *nanos2* and *piwi1* transgenic lines were generated as described previously in (75).

Epigenetic signatures were used to identify putative promoter and enhancer sequences which were then cloned and used to drive the expression of mOrange2 on a p-CRII-TOPO backbone (Schwaiger et al., 2014, Shaner et al., 2004). The plasmids were injected into fertilized eggs at a concentration of 25ng/µl together with I-Sce1 Meganuclease. F0 mosaic transgenic animals were then crossed to wild types to generate F1 heterozygous transgenics.

### RNA probe synthesis

For the generation of in situ hybridization RNA probes, RNA was extracted from mixed embryonic stages using Trizol (ThermoFisher) after the manufacturer’s protocol. cDNA was then synthesized using the Superscript III reverse transcriptase (ThermoFisher) in conjunction with random hexamere and oligo(dT) primers. Primers encompassing the probe sequences were designed using Primer 3 plus (https://www.bioinformatics.nl/cgi-bin/primer3plus/primer3plus.cgi) and primerstats (https://www.bioinformatics.org/sms2/pcr_primer_stats.html). The probe sequences were then amplified by PCR. The PCR products were then run on an agarose gel to confirm the correct product sizes and purified using a PCR purification kit (AnalytikJena). The cloned sequences were then inserted into the pJet1.2 Vector (ThermoFisher), which was then transformed into chemically competent Top 10 E. coli bacteria (Invitrogen). The plasmids were then harvested via miniprep (AnalytikJena), and the desired probe sequences were confirmed using Sanger sequencing (Microsynth). Linearized probe templates were generated by PCR using a T7- or Sp6-promoter containing primer. Probes were then synthesized using the T7 or Sp6 Megascript Kit (Invitrogen) and DIG RNA labeling mix (Roche) (76). The digoxigenin labeled RNA probes were stored in a dilution of 50 ng/µl in a 1:1 mix of formamide and deionized water at -20°C.

### *In situ* hybridization

For embryonic stages, in situ hybridization (ISH) was performed on whole mount 24h gastrula, 2d and 4d planula larvae and 8d old primary polyps. The animals were fixed in 4% PFA/PBS for 1 hour at room temperature, washed three times in PBS with 0.1% Tween 20 (PTW) for 5 minutes each and then transferred into methanol in which they were stored at - 20°C.

ISH was performed after a previously published protocol (Kraus et al., 2016). Minor changes were introduced in the RNA probe detection stap: Anti-Digoxigenin-AP (anti-DIG/AP) Fab fragments (Roche) were used in a dilution of 1:4000 in 0.5% blocking reagent (Roche) in 1x Maleic acid buffer. After incubation overnight at 4°C, the whole-mounts were washed in PTW 10 times for 10 minutes, rinsed twice with alkaline phosphatase buffer (AP-Buffer) and stained with NBC/BCIP dissolved in AP-Buffer. The stained animals were then washed several times in PTW and infiltrated in 86% Glycerol. ISH images were taken using a Nikon 80i compound microscope equipped with a Nikon DS-Fi1 camera.

### Adult tissue in situ hybridization (ISH)

In preparation for ISH, adult polyps were relaxed using 7.5% MgCl2 which was also introduced into the body cavity through the mouth opening using a glass capillary. The animals were then fixed overnight in 4% formaldehyde in PTW on a rotator at 4°C. Afterwards the polyps were dissected into small pieces of ∼5x2mm and washed in 100% methanol. The methanol was regularly replaced until no orange pigment was washed out anymore. Until use, the tissue pieces were stored in methanol at -20°C.

ISH on adult tissues was performed as described previously in (27). In short, PBT (1x PBS with 0.2% Triton X-100) was used instead of PTW in all washing steps. A bleaching step was introduced after rehydration, consisting of an incubation in 0.5% H2O2/5% formamide/0.5X SSC in H2O for 5 minutes. Triethanolamine (TEA) washes were all performed with 1% TEA in PBT. An adapted hybe buffer was used for prehybridization and hybridization step, additionally containing 10% dextrane sulfate (Sigma-Aldrich) and 1% blocking reagent (roche). Prehybridization was extended to overnight and the probe concentration was raised to 0.75 ng/µl. In between the 2x SSC and 0.2x SSC washes, an RNAseT1 incubation (1 U/µl in PBS) for 40 min was introduced to reduce background staining. Lastly, the anti-DIG/AP antibody concentration was raised to 1:2000.

### Generation of *nanos2* knock-out mutant animals

Guide RNAs for CRIPR-mediated mutagenesis (40) were designed using the Chop-Chop online tool (http://chopchop.cbu.uib.no/, (77)) targeting the 3’end of the Nanos-specific zinc-finger RNA-binding domain. The guide RNAs were ordered at IDT-DNA and injected together with Cas9 protein (500ng/µl). Successful mutagenesis was determined by melting curve analysis of a PCR product spanning ∼120 bp around the target site. Mosaic mutant animals were then crossed to wild-type polyps to generate F1 heterozygous mutant families. These were then cross bred to retrieve F2 homozygous knock-out mutants. Mutant animals used in this study have a 497insG mutation in the *nanos2* gene, resulting in a frameshift and premature stop codon.

### Vasa2 immunostaining

For anti-Vasa2 staining, a previously published mouse anti-Vasa2 antibody was used (27), however the immunofluorescence protocol was modified. Adult polyps were starved for 24h and then relaxed using 7.5% MgCl2. They were then transferred into ice-cold fixative consisting of 3.7% formaldehyde/0.4% glyoxal/0.1% methanol/0.2% Triton X-100 in 1x PBS. The fixative was also introduced into the mouth opening using a glass capillary. After 10 min, the animals were dissected into ∼5x2mm pieces in the fixative, and fixation was carried out for an additional 1h at 4°C on a rotator. After fixation, 3 quick washing steps in PBT were performed, followed by 3 permeabilization washes in PBT for 20 min each. The tissue pieces were then transferred into 100% methanol and washed at 4°C with regular replacement of methanol until it remained clear and no pigment dissolved any more. Afterwards a 7 min wash in ice-cold 100% acetone was performed, followed by 2 washes in PBT. The tissue was then blocked for at least 2h at RT in blocking buffer containing 1% BSA/5% sheep serum/20% DMSO/0.2%Triton X-100 in 1x PBS. In parallel, the primary anti-vasa2 AB was pre-absorbed in AB blocking solution with 1% BSA/5% sheep serum/0.1% DMSO/0.2%Triton X-100 in 1x PBS at a concentration of 1:100. The samples were then incubated in AB blocking buffer containing the antibody overnight at 4°C. The samples were then washed 10 times for 10 min each in PBT to remove the primary AB. The secondary antibody (goat anti-mouse 488, Sigma-Aldrich) was then preabsorbed in AB blocking buffer and the samples were blocked in the same solution, before being incubated in the secondary AB containing solution overnight at 4°C. Another 10 washing steps for 10 min in PBT were performed and the tissue pieces were then gelatine sectioned and stained with 5µg/ml DAPI. The sections were subsequently mounted onto slides and imaged using a confocal microscope (Leica Sp8).

### EdU staining

Transgenic animals were incubated in 20µM EdU solution in 2% DMSO in NM for 5h for single cell dissociations and for 24h when used for gelatine sections. For single cell spreads, the animals were then dissociated using the Collagenase/Papain protocol and adhered to superfrost slides as described above. After fixation a permeabilization step using 0.5% Triton X-100 in PBS was performed for 30 min, followed by 3 washing steps in PBS and the EdU detection reaction as per the manufacturer’s protocol for 2 min on the slides. For additional immunostaining, the cells were washed 2 times in PBT and subsequently incubated in blocking solution consisting of 1% BSA and 20% sheep serum in PBT for 2h in a sealed humid chamber. The primary antibody was also blocked for 2h and then slides were incubated with the AB solution overnight. To wash off the primary AB, five washing steps with PBT were performed. The slides and secondary antibody were then again incubated in blocking solution for 2h and the secondary AB was hybridized overnight. The secondary AB was once again removed through five washing steps in PBT. Before mounting, the cells were rinsed with PBS which was subsequently replaced by Vectashield for imaging. For gelatine sectioning, the animals were washed several times with NM after the EdU incubation, and relaxed using 7.5% MgCl2 in NM, diluted 1:10. They were then fixed for 16h in 4% PFA in PBS at 4°C. The animals were then cut into smaller pieces of ∼5x2mm and washed for several hours in 100% methanol, regularly replacing the methanol until no more pigment was washed out. The tissue pieces were then rehydrated in PBS and put into 10% gelatine in PBS to infiltrate for 1-2h. The gelatine was then poured into molds, which were hardened at 4°C and re-fixed overnight in 4% PFA at 4°C. The gelatine plugs containing the tissue pieces were then rinsed in PBS again and 50µM sections were prepared using a vibratome (Leica). The sections were then stored in PBS until they were incubated in DAPI and phalloidin and subsequently mounted.

## Acknowledgements

The authors thank all fellow members of the Technau group for constructive discussions and the Core Facility Cell Imaging of the Faculty of Life Sciences (CIUS, University of Vienna) for facilitating confocal imaging. We thank our facility staff members Angela Caballero Alfonso, Vendula Stejskalova and Wolfgang Göschl for animal husbandry. Further we thank Daniela Praher for providing adult gonad *piwi1* ISH pictures and Paul Knabl for the Vasa2 Antibody immunostaining protocol. This work was funded by the Austrian Science Fund to UT (P27353) and AGC (P31018).

## Supporting Information

**S1 Fig: *nanos2* and *piwi1* expression in developmental stages.** Numbers indicate the developmental stage in each panel, 1: 24h gastrula stage, 2: 3d planula stage, 3: 7d primary polyp; A: ISH visualizing *nanos2* expression throughout development; B: *nanos2*::mOr expression; C: ISH visualizing *piwi1* expression throughout development; D: *piwi1*::mOr expression; E-F: *nanos2*and *piwi1* reporter gene constructs.

**S2 Fig: Phylogenetic analysis of Nanos proteins.** Bayesian consensus tree of metazoan nanos proteins; Numbers at the branches indicate bootstrap support (1000 iterations). *nanos* genes have duplicated in the cnidarian lineage and triplicated at the base of the teleost lineage independently. Dme: *Drosophila melanogaster*; Hro: *Hirundo robusta*; Dre: *Danio rerio*; Hsa: *Homo sapiens*; Mmu: *Mus musculus*; Xla: *Xenopus laevis*; Bfl: *Branchiostoma floridae*; Hvu: *Hydra vulgaris*; Che: *Clytia hemisphaerica*; Hec: *Hydractinia echinata*; Pca: *Podocoryne carnea*; Nve: *Nematostella vectensis*; Emu: *Ephydatia muelleri*.

**S3 Fig: Long term EdU label-retention assay in *piwi1*::mOr transgenic animals. A-C**: EdU signal is retained in few mOr+ cells in the mesentery epithelia over 1 month, marking a subpopulation of slowly cycling cells. **A, B**: 1 month label retention in mesenteries. **A**’, **B**’: DAPI only. **A**’’, **B**’’: EdU only. **A**’’’, **B**’’’: mOrange only.

**S4 Fig: *nanos2* genomic coding sequence of *nanos2* KO mutants.** Nanos Zinc-finger is highlighted in turquoise, 1bp insertion is highlighted in red.

