## Supplementary figures and images for "*Nanos2+* cells give rise to germline and somatic lineages in the sea anemone *Nematostella vectensis*" (from another version of this preprint: version 1, 2023-12-08)

### Supplemental Figure 1

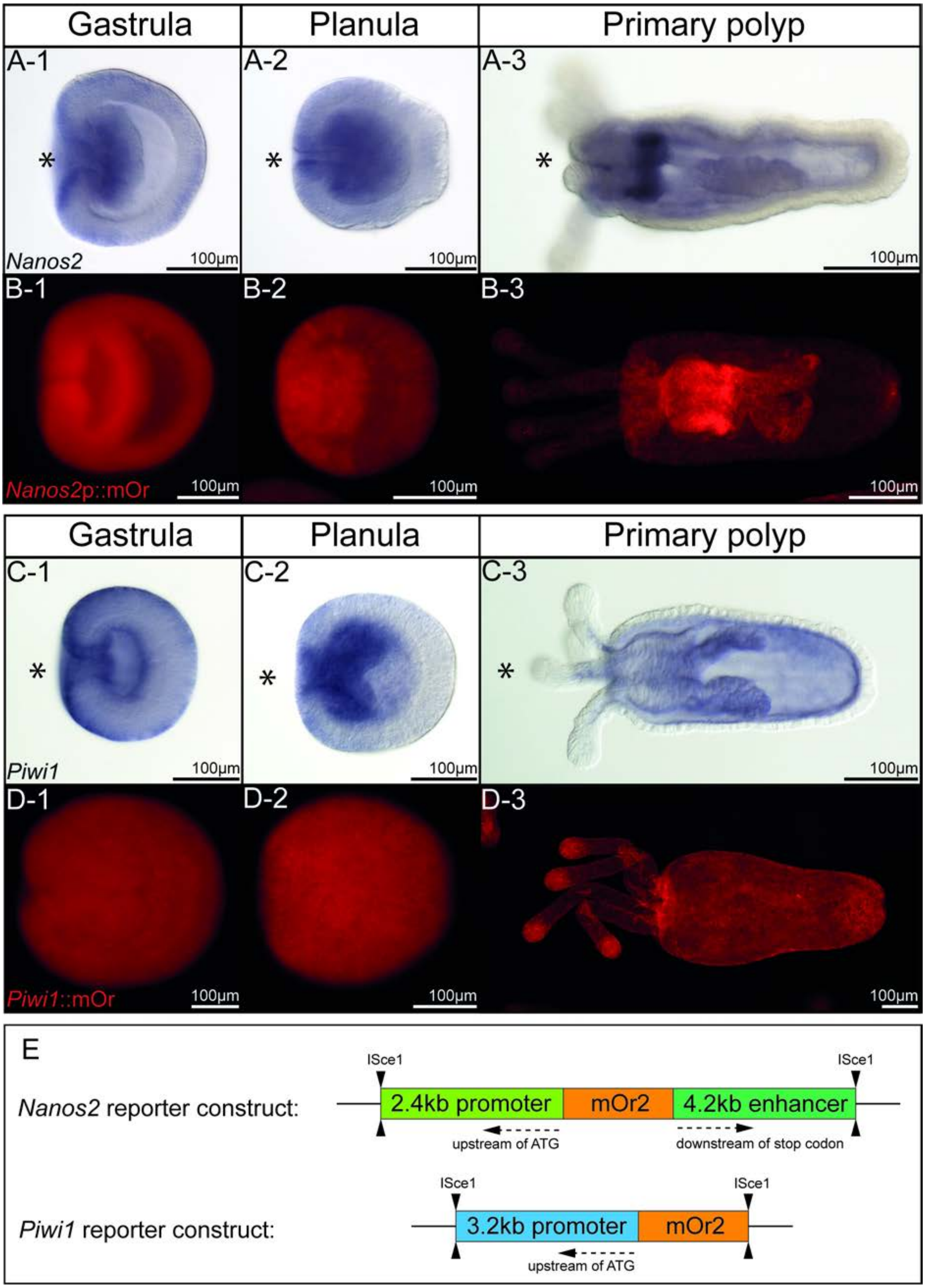

### Supplemental Figure 2

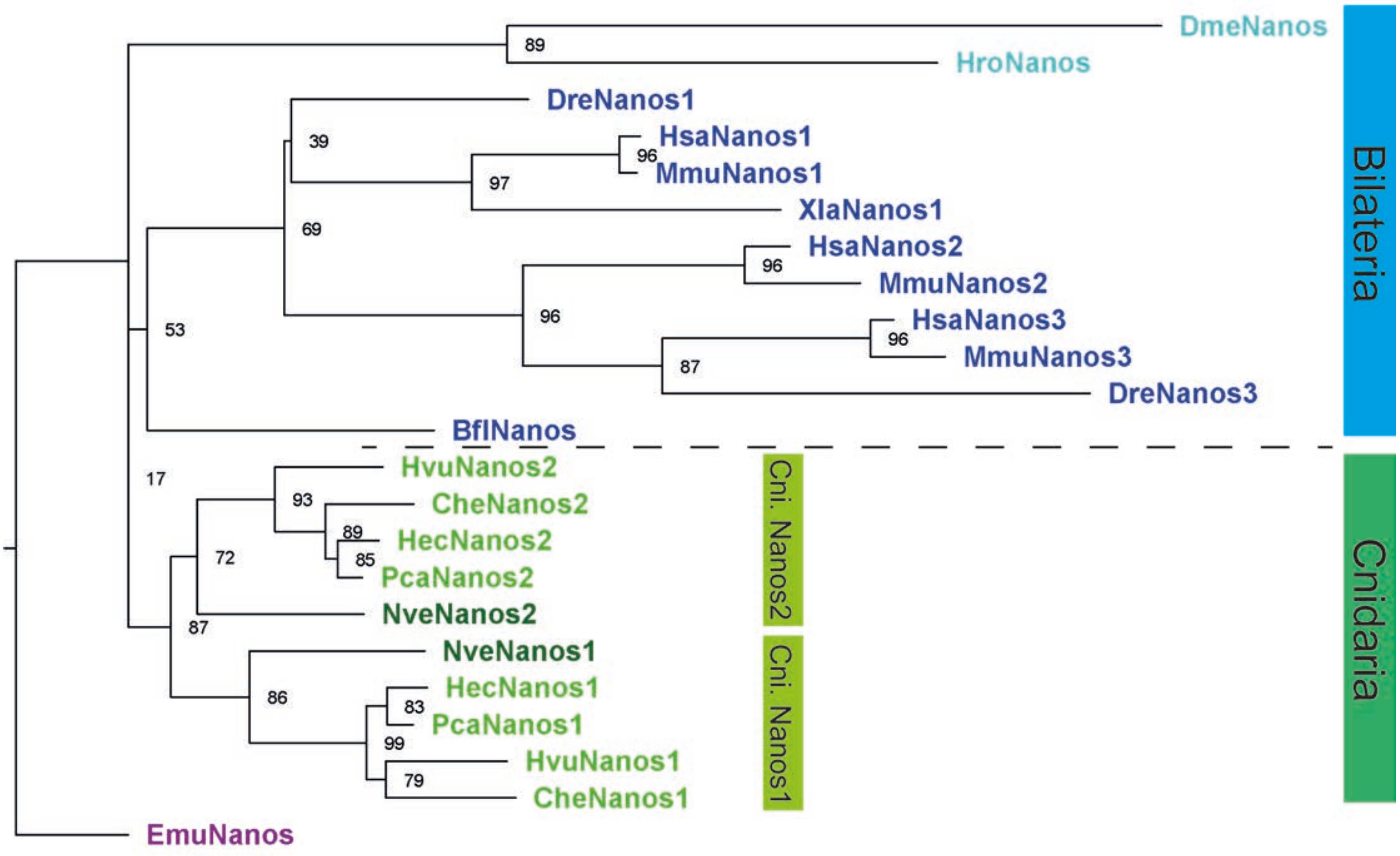

### Supplemental Figure 3

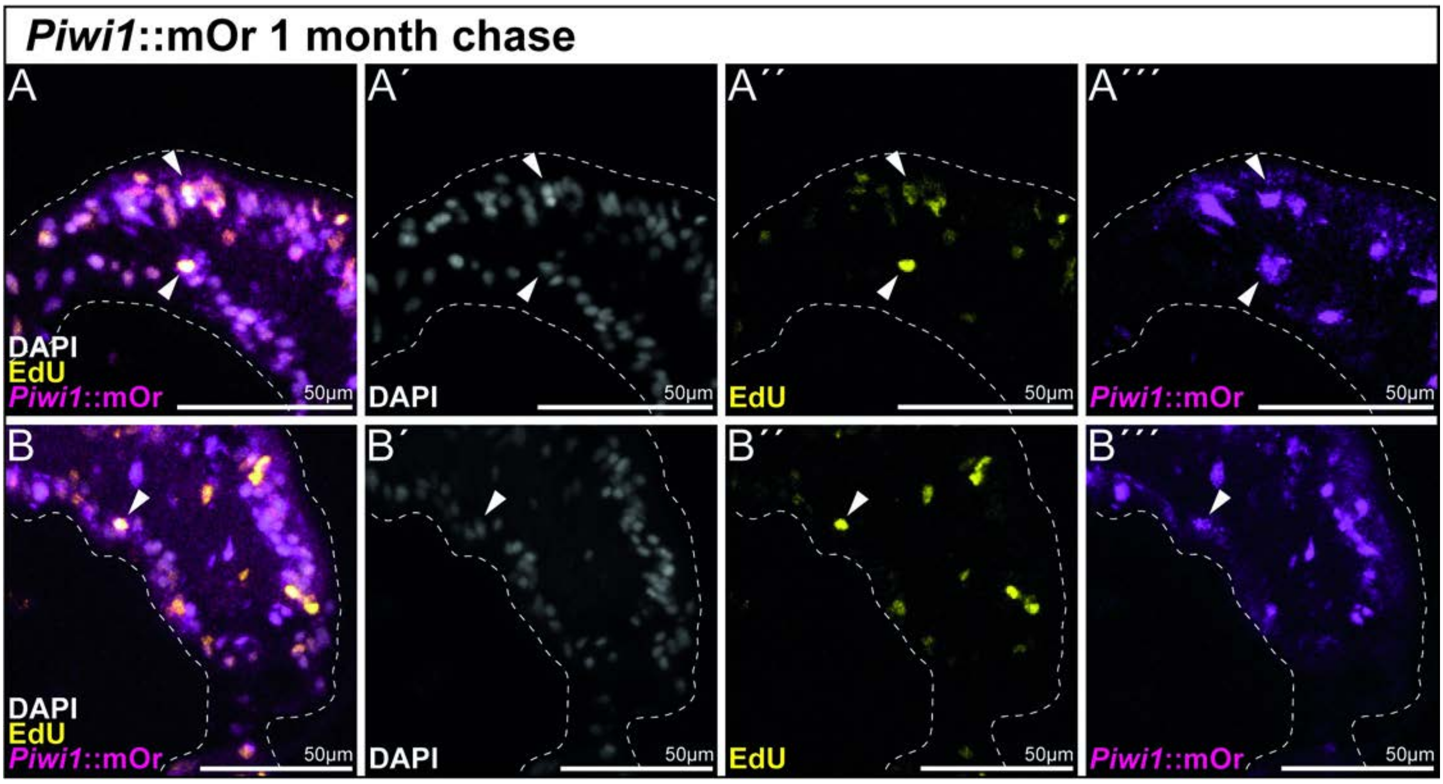
