## Supplemental Figure 4 for "*Nanos2+* cells give rise to germline and somatic lineages in the sea anemone *Nematostella vectensis*" (from another version of this preprint: version 1, 2023-12-08)

```
ATGTCTTTGAGTGAGGTGATGTTAAAAACGCTGATTCCCAGCAACTTTTTGGACTCAGACGACGAAGAGATAAAG
TCGTTAACGTGCACTACGGAGAGCGATGATTTGCAGAAAAATCAGTATACCAACTACAACATGTTCAACGACTAC
TTCGGGCTCAGTAATCTGGTGCAAAAAGTGCGGCTCGCTAATAATGATTTCCCGTATCTCGGCAGCGATGGCTAC
GGCGGCAAACTGCACCGTCGTGACCGATTTGACAGCATTGATTCGGAGAACTCGTCTTCTGGAAGCTGTGATTCC
TTCACGGATCACGGCCTGACATTCGGCGAGGGATCATTTCAGCCATTTAGCACATCATCTATCCTGCACAGTGCA
GGGAAAAGGAAAGAAGTTGCGTTGCCGCCAAGTCGAAGGCTCGGAAAGCCGACCGCGAAGCTCAGCTCCCGGT
GCAAATCGTCAAGTGTGCGTGTTCTGCAGAAACAACGGTGAGAGCGGAGGAGGTGTACGCTTCACATGTGTTAAA
AAGTGCCGATGGGAAAACTACTTGCCCGATCCTGAGAGCCTACACTTGTCCGATTTGCAAGGCAAGCGGCGATGA
TTCGCATACGATAAAATATTGCCCGCAAAATCAGCAAACGCAAGGAAATGGCCAGCTTCCTCCTCCACCAGTCAA
ACCACCAACCAGCACAGCACAGCCAATCGCTCGCTCCACGCGGGGAATAATGCCGCCAAAACGCCGCTAGACTAA
```
